# Complete mitochondrial genomes of Thai and Lao populations indicate an ancient origin of Austroasiatic groups and demic diffusion in the spread of Tai-Kadai languages

**DOI:** 10.1101/063172

**Authors:** Wibhu Kutanan, Jatupol Kampuansai, Metawee Srikummool, Daoroong Kangwanpong, Silvia Ghirotto, Andrea Brunelli, Mark Stoneking

**Author notes:** Corresponding authors; 1. Professor Dr. Mark Stoneking, Department of Evolutionary Genetics, Max Planck Institute for Evolutionary Anthropology Deutscher Platz 6, D04103 Leipzig, Germany. Dr. Wibhu Kutanan, Department of Biology, Faculty of Science, Khon Kaen University, Mittapap Road, Khon Kaen, 40002, Thailand.

## Abstract

The Tai-Kadai (TK) language family is thought to have originated in southern China and spread to Thailand and Laos, but it is not clear if TK languages spread by demic diffusion (i.e., a migration of people from southern China) or by cultural diffusion, with native Austroasiatic (AA) speakers switching to TK languages. To address this and other questions, we obtained 1,234 complete mtDNA genome sequences from 51 TK and AA groups from Thailand and Laos. We find high genetic heterogeneity, with 212 haplogroups. TK groups are more genetically homogeneous than AA groups, with the latter exhibiting more ancient/basal mtDNA lineages, and showing more drift effects. Modeling of demic diffusion, cultural diffusion, and admixture scenarios consistently supports the spread of TK languages by demic diffusion. Surprisingly, there is significant genetic differentiation within ethnolinguistic groups, calling into question the common assumption that there is genetic homogeneity within ethnolinguistic groups.

---

Thailand and Laos are regarded as the geographical heart of Mainland Southeast Asia (MSEA) (Fig. 1). Archaeological evidence suggests a long history of human occupation of the area, with the oldest human remains dated to 46-63 thousand years ago (kya) from Tam Pa Ling Cave^1^, and cultural remains dating to 35-40 kya^2–3^. A potential role for Thailand/Laos as a corridor between southern China and Island Southeast Asia (ISEA) is further indicated by archaeological evidence for agricultural communities that may have expanded from the center of the Yangtze valley during the Neolithic period^4–5^.

**Figure 1.**
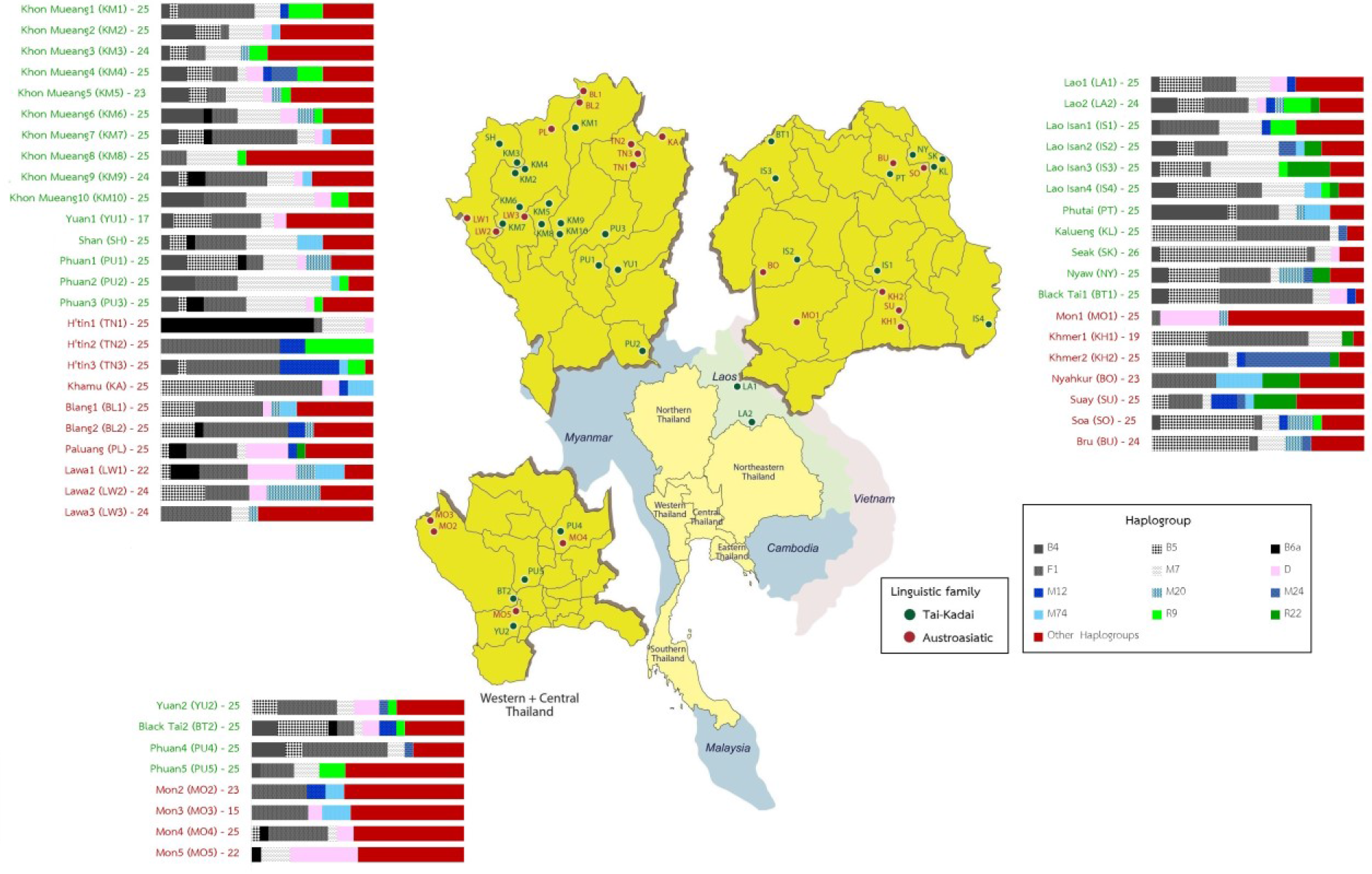
Map showing the geographic locations of the studied populations and their language family affiliation. Bar plots illustrate the relative frequency of major haplogroups by population. Dark and white shades show haplogroups B, F and M7, which are specific to Southeast Asian populations, whereas the remaining haplogroups (D, M12, M20, M24, M74, R9, R22 and other haplogroups) are represented by various colors.

There is also considerable linguistic diversity, with five language families (Tai-Kadai (TK), Austroasiatic (AA), Sino-Tibetan (ST), Hmong-Mien (HM) and Austronesian (AN)), spoken in the area. Most people speak TK languages (94.40%, in Thailand and 69.60% in Laos) while AA is the second most common language family (4.10% in Thailand and 22.70% in Laos)^6^. However, the AA family is more diverse (27 languages in Thailand and 47 languages in Laos) than TK (16 languages in Thailand and 21 languages in Laos). The ST and HM families are concentrated in the area of northern and northwestern Thailand as well as northern and central Laos (ST: 19 languages in Thailand and 11 languages in Laos; HM: 3 languages in Thailand and 4 languages in Laos). The AN family is restricted to southern Thailand with just 6 languages^6^. Both major families (AA and TK) are widespread across Asia; there are 167 AA languages spoken by ~102 million people from South Asia (Bangladesh and India) to southern China and MSEA, including Malaysia; and 92 TK languages spoken by ~80 million people in northeast India, southern China, Vietnam, Myanmar, Cambodia, Thailand and Laos^6^. Although the origin and spread of AA is debatable^7–8^, AA people are generally considered to be descended from the earliest inhabitants of the region^9–10^. TK is generally considered to have arisen in southeast China prior to 2.5 kya and then spread to SEA between 1-2 kya^11–12^.

Although archaeological and linguistic evidence point to an expansion from southern China, physical anthropological studies indicate that the present-day Thai people resemble ancient people^13^ as well as modern AA people in northern Thailand^14^. Therefore, there are two competing hypotheses concerning the origin of the modern Thai/Lao TK people: (1) a demic expansion of people from southern China that brought their genes, culture, and language to Thailand/Laos; or (2) a cultural diffusion from southern China that resulted in native AA peoples adopting the TK language and culture. This general question of demic vs. cultural diffusion is a longstanding one concerning expansions in other parts of the world, particularly those involving languages and/or agricultural practices, e.g. expansions associated with Indo-European, Bantu, Han and Austronesian languages^15–22^. While genetic studies have proven informative in distinguishing between demic vs. cultural diffusion in these other contexts, to date genetic studies have not been applied to this question with respect to TK peoples. In particular, previous mitochondrial (mt) DNA studies on Thai/Lao populations were too limited to address this question via phylogenetic or simulation based analyses^23–25^. Therfore, in order to address the role of demic vs. cultural diffusion in the origins of the TK people as well as investigate other aspects of Thai/Lao prehistory, we analyze here 1,234 complete mtDNA genome sequences from 51 Thai/Laos populations, comprising a comprehensive sampling of TK and AA genetic diversity

## Results

### Genetic diversity is higher in TK than in AA groups

For the 1,234 mtDNA genome sequences obtained, there are 761 distinct sequences (haplotypes) belonging to 212 haplogroups (Supplementary Table 1). The summary statistics for the genetic diversity in each population are provided in Supplementary Table 2. Haplotype diversity (*h*) ranges from 1.00 in the LA2 (see Fig. 1 for population locations and population abbreviations) to 0.80 in the TN2 group. The SK, BO and TN1 groups also exhibit *h* values somewhat lower than the remaining populations; the same trend is observed for haplogroup diversity, as relatively large values are observed in almost all populations except in TN1, TN2, SK and BO. Both nucleotide diversity (*π*) and mean number of pairwise differences (MPD) are also the lowest in the TN1 group (0.0013 and 21.41, respectively), while the largest values are observed in the MO2 group (0.0026 and 42.6, respectively).

Haplotype and haplogroup diversity values as well as the number of segregating sites are significantly higher for TK than for AA groups (Mann-Whitney U tests: *h*: Z = 3.34, *P* = 0.0008, haplogroup diversity: *Z* = 3.53, *P* = 0.0004, number of segregating site: *Z* = 2.85, *P* = 0.0044). However, the *π* values of AA groups are not significantly differ from those of the TK groups (*Z* = 1.45, *P* = 0.15).

### Greater genetic heterogeneity of AA groups

The multidimensional scaling (MDS) analysis (Fig. 2a-b) revealed that in the third dimension AA and TK groups tended to be separated; this separation was more apparent when three outliers were excluded (Fig. 2c-d). The correspondence analysis (CA) analysis based on haplogroup frequencies (Supplementary Fig. 1) indicates that specific haplogroups are associated with the populations showing relatively high levels of genetic differentiation, namely: haplogroup B6a in TN1; haplogroup M12a1a in TN3; haplogroup F1a1a in TN2 and BO; and haplogroup B5a1d in SK and KA. Overall, the MDS and CA analyses revealed greater genetic heterogeneity among AA than TK groups. This result is supported by the analysis of molecular variance (AMOVA) (Table 1), as 11.44% of the variance is among AA populations, compared to 4.74% for the TK populations. However, neither linguistic nor geographic classifications of the populations provide a good match to the underlying genetic structure of the Thai/Laos populations, as in all such classifications the among-population component of the variance is higher than the among-group component (Table 1). Moreover, the Mantel test for the correspondence between genetic and geographic distances between populations is not significant in all types of geographic distances tested (great circle distance: *r* = 0.03, *P*= 0.31, least cost path distance: *r* = 0.04, *P* = 0.30 and resistance distance: *r* = −0.65, *P* = 0.75). Thus, the genetic structure of the Thai/Laos populations is more complicated than would be predicted from either linguistics or geography.

**Figure 2.**
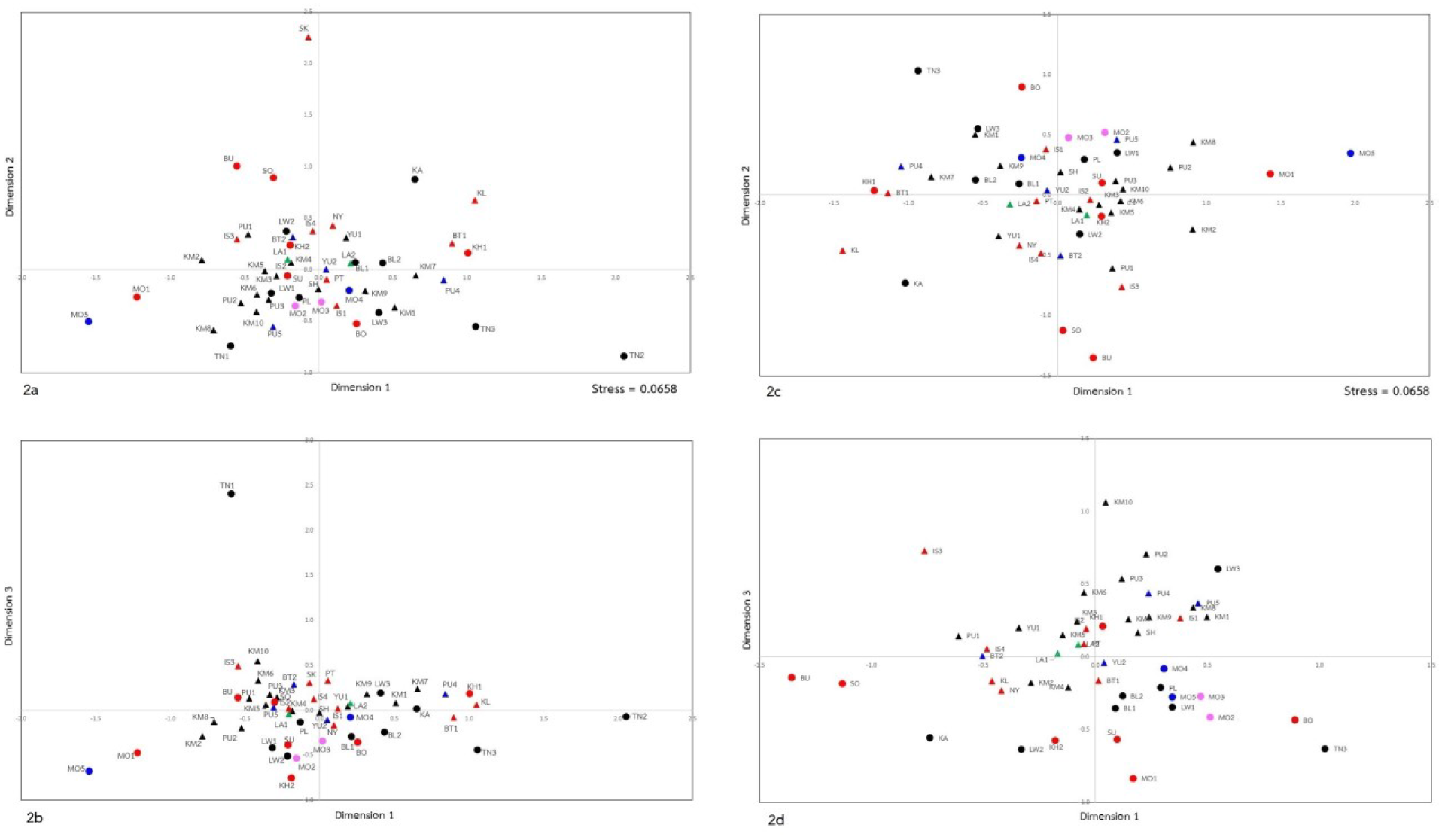
MDS plot of dimension 1 vs. dimension 2 (2a and 2c) and dimension 1 vs. dimension 3 (2b and 2d) based on the *Φ*_*st*_ genetic distance matrix among the entire set of 51 populations (2a and 2b) and after removal of three outliers, namely TN1, TN2 and SK (Fig. 2c and 2d). Population abbreviations are provided in Fig. 1. Triangles and circles represent TK and AA speaking populations, respectively. Black, red, dark blue and pink colors indicate North, Northeastern, Central and West geographic regions of Thailand respectively; green indicates the two Lao populations.

**Table 1.** Analysis of Molecular Variance (AMOVA) results.

| Grouping | Number of groups | Percent variation |  |  |
| --- | --- | --- | --- | --- |
|  |  | Among groups | Among population (within group) | Within population |
| Geography |  |  |  |  |
| Geography 1 | 5 | 0.07 | 7.63** | 92.3** |
| Geography 2 | 4 | 0.36 | 7.77** | 91.86** |
| Northern Thailand | 1 | - | 7.76** | 92.24 |
| Northeastern Thailand | 1 | - | 8.69** | 91.31 |
| Central Thailand | 1 | - | 6.83** | 93.17 |
| Western Thailand | 1 | - | -0.43 | 100.43 |
| Laos | 1 | - | 0.66** | 99.34 |
| Language |  |  |  |  |
| Language 1 | 2 | 0.49* | 7.42** | 92.1** |
| Language 2 | 6 | 2.56** | 6.01** | 91.43** |
| Language 3 | 10 | 2.42** | 5.68** | 91.9** |
| Austroasiatic | 1 | - | 11.44** | 88.56 |
| Tai-Kadai | 1 | - | 4.74** | 95.26 |
| Ethnicity |  |  |  |  |
| Mon | 1 | - | 7.1** | 92.9 |
| H'tin | 1 | - | 25.71** | 74.29 |
| Lawa | 1 | - | 7.78** | 92.22 |
| Khmer | 1 | - | 11.10** | 88.90 |
| Khon Mueang | 1 | - | 3.43** | 96.57 |
| Lao Isan | 1 | - | 2.31** | 97.69 |
| Phuan | 1 | - | 5.29** | 94.71 |

\*significant at 0.05 level; \*\* significant at 0.01 level.
Geography 1: Northern Thailand, Northeastern Thailand, Central Thailand, Western Thailand, Laos)
Geography 2: (Northern Thailand, Northeastern Thailand, Central Thailand, Western Thailand)
Language 1: (Austroasiatic, Tai-Kadai)
Language 2: (Northern Tai, Southwestern Tai, Monic, Southern Monic, Eastern Mon-Khmer, Northern Mon-Khmer)
Language 3: (Northern Tai, Chiang Saen, Lao-Phutai, Northwestern Tai, Monic, Southern Monic, Palaungic, Khmuic, Khmer, Katuic)

Greater genetic homogeneity among the TK populations was also reflected in the haplotype sharing analysis (Supplementary Table 3), which showed that they shared more haplotypes than the AA populations. In particular, the various KM populations shared a number of haplotypes, as did the PU populations, indicating some recent genetic exchange/ancestry among populations within the same ethnolinguistic group. The highest number of shared haplotypes is five, which are shared among the KM5-KM6 and PU2-PU4 groups. Many haplotypes in the PU are shared with almost all of the other TK populations. Among the AA populations, despite the relatively large genetic differences between the TN2 and TN3 populations, they share four haplotypes. Overall, only four populations (IS3, SK, MO1 and MO4) did not share any haplotypes with any other population.

### Significant genetic differentiation within ethnolinguistic groups

Surprisingly, we observed striking and significant genetic differences between populations classified as the same ethnolinguistically but sampled from different locations. This can be seen in the MDS analysis (Fig. 2a-b), in which two of the three most extreme outliers are from the same ethnolinguistic group, namely two of the three AA-speaking H’tin groups, TN1 and TN2 (the third outlier is the SK, a TK-speaking group from northeastern Thailand). In fact, the MDS analysis shows that in many cases populations from the same ethnolinguistic group are not genetically similar. This is further indicated by an AMOVA for each separate ethnolinguistic group that was sampled from multiple locations (Table 1); in all such instances, the among-populations variance component is significantly different from zero. This unexpected high degree of heterogeneity within the same ethnolinguistic group contributes to the lack of correspondence between the genetic structure of the Thai/Laos populations and their geographic/linguistic relationships.

### Relationships with other Asian populations

The genetic relationships of 113 Asian populations (51 from the current study and 62 from the literature; Supplementary Table 4) as revealed by MDS analysis indicated, in general, population clustering by both language family and macro-geographic scale (Fig. 3). The SEA populations who speak AN, AA and TK languages are largely separated from North and South Asian populations. The AN and AA groups are further differentiated by the second dimension with the intermediate position of the TK populations among them. These results are also seen in the Neighbor Joining (NJ) tree, with the East Asian populations separated from the North and South Asian populations (Supplementary Fig. 2). Most of the AN groups from Taiwan, Philippines, and Island Southeast Asia (ISEA) are separated from the Thailand TK and AA populations. The TK and AA populations are mostly intermingled with a few AN populations also clustering with them. Overall, TK and AA populations are closed to AN population in both MDS (Fig. 3) and NJ tree (Supplementary Fig. 2). Among the presently studied populations, again, the TN1, TN2 and SK are extremely divergent (in keeping with their relatively low amounts of genetic diversity) but they nonetheless cluster with their neighbors from Thailand. There is also a clear division in the AA populations: MO1 and MO5 show affinities with populations from Myanmar and India, reflecting their genetic relatedness (Fig. 3), and are distinct from the other Mon and the other Thai populations. This could reflect either common ancestry of MO1 and MO5 with groups from Myanmar and India and/or gene flow. Surprisingly, even though the two Khmer populations (KH1 and KH2) from northeastern Thailand have close geographic proximity and shared haplotypes, they are genetically distinct from one another and from an ethnolinguistically-related group, the Cambodian Khmer (KH_C).

**Figure 3.**
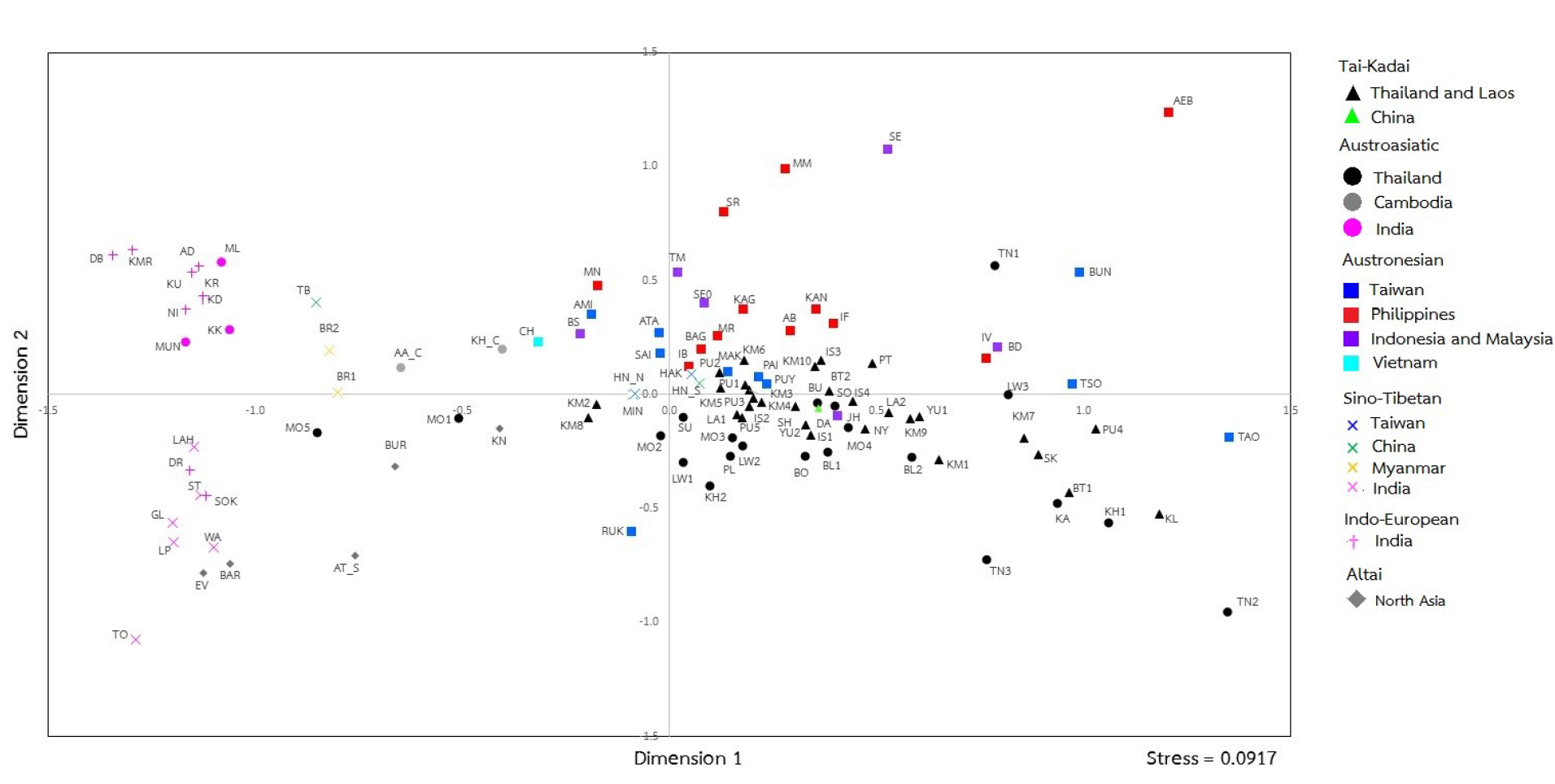
MDS plot based on *Φ*_*st*_ genetic distance matrix from mtDNA genomes among the presently studied populations and other populations from the literature. Population abbreviations are provided in Fig. 1 and Supplementary Table 4.

### mtDNA lineages

The above population relationships are based on analyses of the entire set of mtDNA sequences; additional insights come from considering the distribution and other characteristics of specific haplogroups. Among the 1,234 mtDNA genomes belonging to 212 haplogroups, F1 is by far the predominant lineage (21.80%), followed by B5 (13.13%), M7 (11.02%) and B4 (6.00%) (Fig. 1). All of these haplogroups are common in SEA populations and predominate in most of the studied populations, with the exception of two TK (KM8 and PU5) and 12 AA (PL, LW1-LW3, KH2, BO, SU and MO1-MO5) populations (Fig. 1). Haplogroup coalescent times using Bayesian Markov Chain Monte Carlo (MCMC) estimates (BE) and credible intervals (CI) by haplogroup are shown in Fig. 4. A schematic phylogeny of the main haplogroups, based on Bayesian MCMC analyses, is provided in Fig. 5, while full Bayesian maximum clade credibility (MCC) trees by haplogroup are presented in Supplementary Fig. 3. Networks of the sequences in each haplogroup are presented in Supplementary Fig. 4, and frequency maps of some haplogroups are in Supplementary Fig. 5. A detailed discussion of each main haplogroup is in the Supplementary Text; here we summarize the main findings.

**Figure 4.**
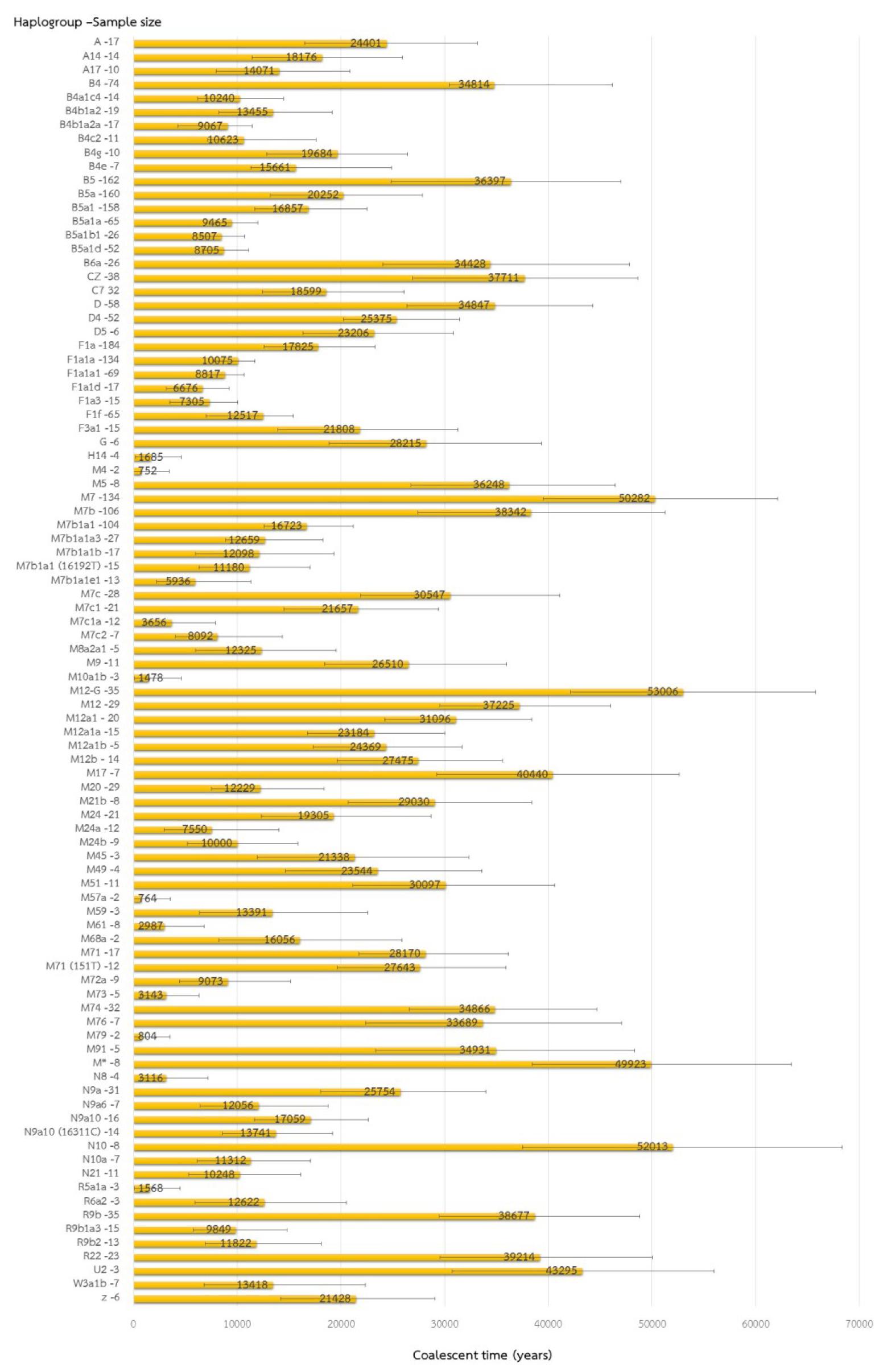
The Bayesian estimates (BE) of coalescent times with 95% credible intervals (CI) for each haplogroup.

**Figure 5.**
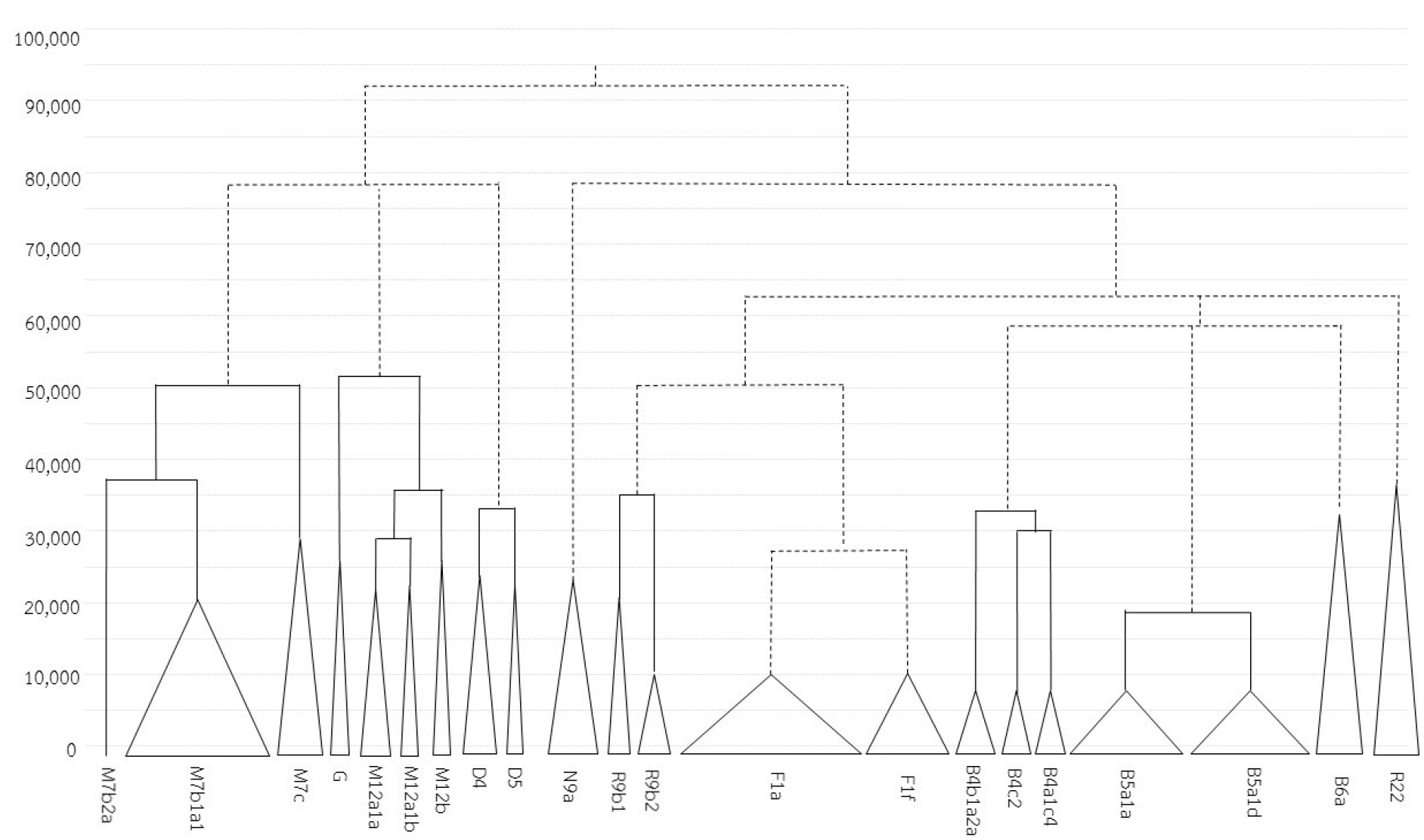
The Schematic Bayesian MCMC tree of the major haplogroups found in this study. Bayesian maximum clade credibility trees were constructed for each haplogroup with parameters as described in the Methods and then manually combined (dashed lines) based on PhyloTree mtDNA tree build 17. The full Bayesian maximum clade credibility tree for each haplogroup is shown in Supplementary Fig. 3.

The haplogroup profiles by population emphasize the greater genetic heterogeneity in AA groups than in TK groups (Fig. 1 and Supplementary Table 1). Some AA groups have extremely high frequencies of particular haplogroups, indicating the pronounced effect of genetic drift; examples include: R9b2 with a frequency of 32.00% in TN2; R22 with frequencies of 17.39% in BO and 20.00% in SU; D4 with frequencies of 28.00% in MO1, 31.81% in MO5, 22.73% in LW1, and 20.00% in PL; and B6a with a frequency of 72.00% in TN1. Overall, the greater heterogeneity in haplogroup distribution and pronounced haplogroup frequency differences are consistent with an older presence of AA groups in Thailand.

Some haplogroups prevalent in South Asia also occur in some AA groups, especially the Mon groups. These include D4, mentioned above, as well as W3a1b, which is reported here for the first time in MSEA. W3a1b was found in two Mon populations (24.00% in MO1 and 4.35% in MO2); these haplogroups provide further evidence for genetic connections between these Mon groups and South Asia.

Although many haplogroups are shared between MSEA and ISEA, there are distinct differences in the distribution of some sublineages. For example, haplogroup B4 is widespread throughout SEA; in our study it is almost entirely restricted to TK groups (Fig. 1 and Supplementary Table 1) where it occurs as three primary sublineages, namely B4b1a2a, B4a1c4 and B4c2, all of which have been reported previously in MSEA^21,26^. Several other B4 sublineages characteristic of Taiwan (e.g., B4b1a2h, B4b1a2f and B4b1a2g^27^), the Philippines (e.g., B4b1a2b, B4b1a2c and B4b1a2d^28^) and Oceania (e.g., B4a1a1a^29^) were not found in our study, in agreement with previous studies^26,30^. Overall, the lack of sharing of recent sublineages indicates a lack of recent contact between MSEA and ISEA (Supplementary Fig. 4).

Finally, the more extensive sampling of Thai/Laos mtDNA sequences in this study has resulted in much deeper ages for some haplogroups that were poorly sampled in previous studies. For example, we estimate that haplogroups R9b and R22 both coalesce at ~39 kya (Fig. 4), compared to previous estimates of ~29 kya^31^ and ~19 kya^26^ respectively. Moreover, while R9b and R22 have been suggested to originate in southern China^31^ and ISEA^26,32^ respectively, northeastern Thailand is also a potential source for these haplogroups (Supplementary Fig. 5).

### Population size change trends over time

The Bayesian Skyline Plots (BSP) in each of the 51 populations individually (Supplementary Fig. 6) reveal four overall trends in change in *N_e_* over time (Fig. 6). The most common trend (observed in 24 TK and 13 AA groups) is an increase in *N_e_* around 50 to 40 kya, followed by stability and then a decline around 2 kya (Fig. 6a). A different trend is observed in most of the ethnic Lao populations (IS and LA) and one KM population; the IS1, IS2, LA2 and KM5 populations expanded continuously but stay stable for the present time (Fig. 7b) while IS4 and LA1 show population expansions at around 50 kya and again around 10 kya (Fig. 6c). Another pattern of observed demographic change (Fig. 6d) is a stable *N_e_* since the upper Paleolithic and then a sudden decline during the last 2 kya, which could produce a larger drift effect, and is seen in 8 AA groups.

**Figure 6.**
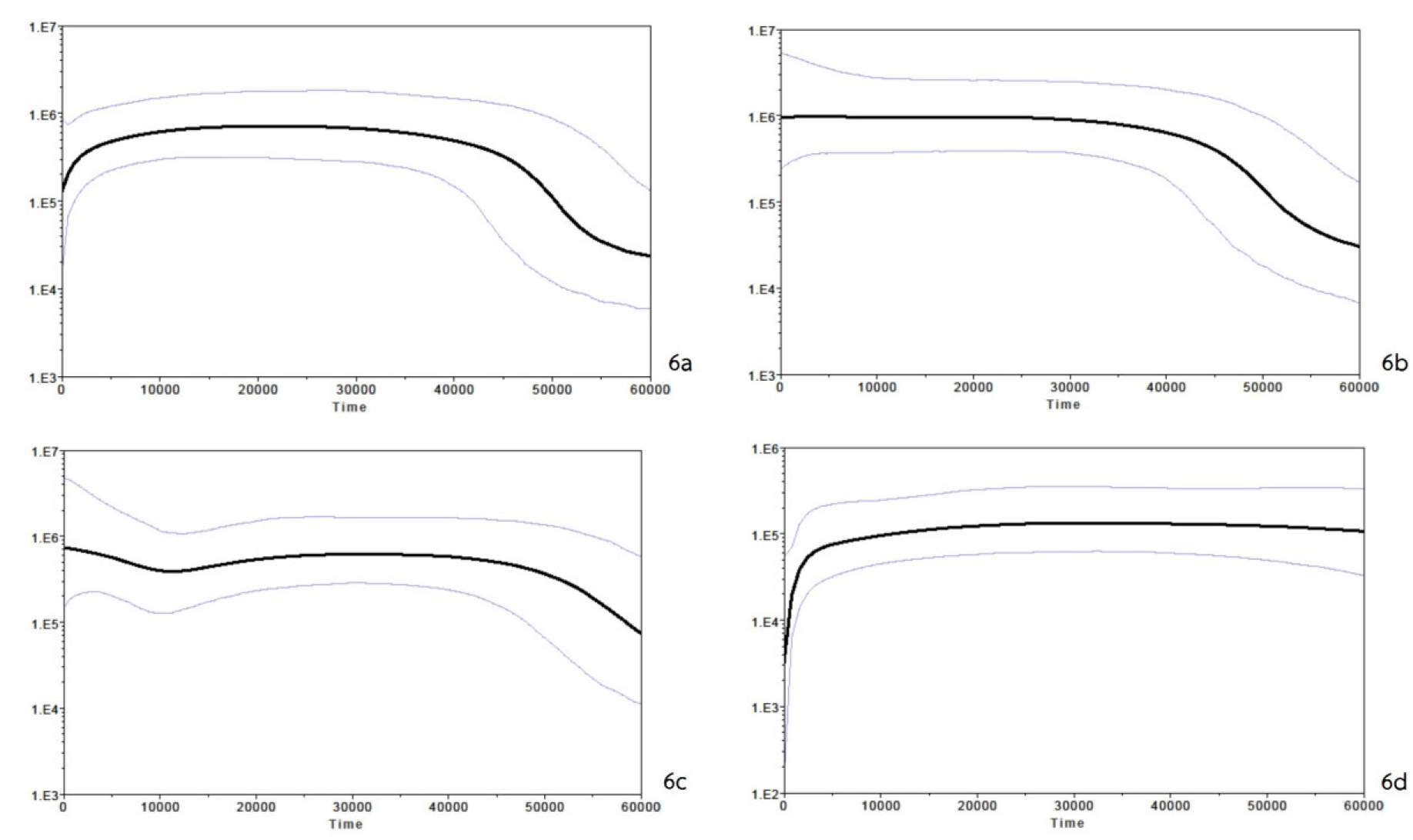
Four different trends in fluctuation in maternal effective population size (y-axis) through time from the present in unit of years (x-axis) observed in the individual Bayesian skyline plots (BSP) for the 51 populations (Supplementary Fig. 6). The median estimate and the 95% highest posterior density limits are indicated by thick and thin lines, respectively. The plots were generated with 10,000,000 chains with the first 1,000,000 generations discarded as burn-in. Most populations (KM1-KM4, KM6-KM10, YU1-YU2, SH, IS3, PT, NY, KL, SK, BT1-BT2, PU1-PU5, MO1-MO5, KH2, BU, SO, SU, LW1, PL, BL1-BL2) show the trend in 6a; KM5, IS1-IS2 and LA2 show the trend in 6b; IS4 and LA1 show the trend in 6c; and KH1, BO, TN1-TN3, KA and LW2-LW3 show the trend in 6d.

### Testing models of demic diffusion vs. cultural diffusion vs. admixture

To address the role of demic vs. cultural diffusion in the origins of Thai/Lao people, we proposed and tested demographic models according to immigrant vs. indigenous hypotheses (Fig. 7). The immigrant hypothesis (or demic diffusion) states that the nowadays TK people descend primarily from the TK-speaking groups from southern China who migrated southward in the last 1 to 2 kya^11–12^. By contrast, the indigenous hypothesis (or cultural diffusion) suggests that the TK people descend primarily from native AA inhabitants who shifted culturally and linguistically^9^. In addition, we consider another possible scenario, namely admixture, which explains the dual origin of the current TK people as reflecting a genetic mixing of incoming TK and indigenous AA groups.

**Figure 7.**
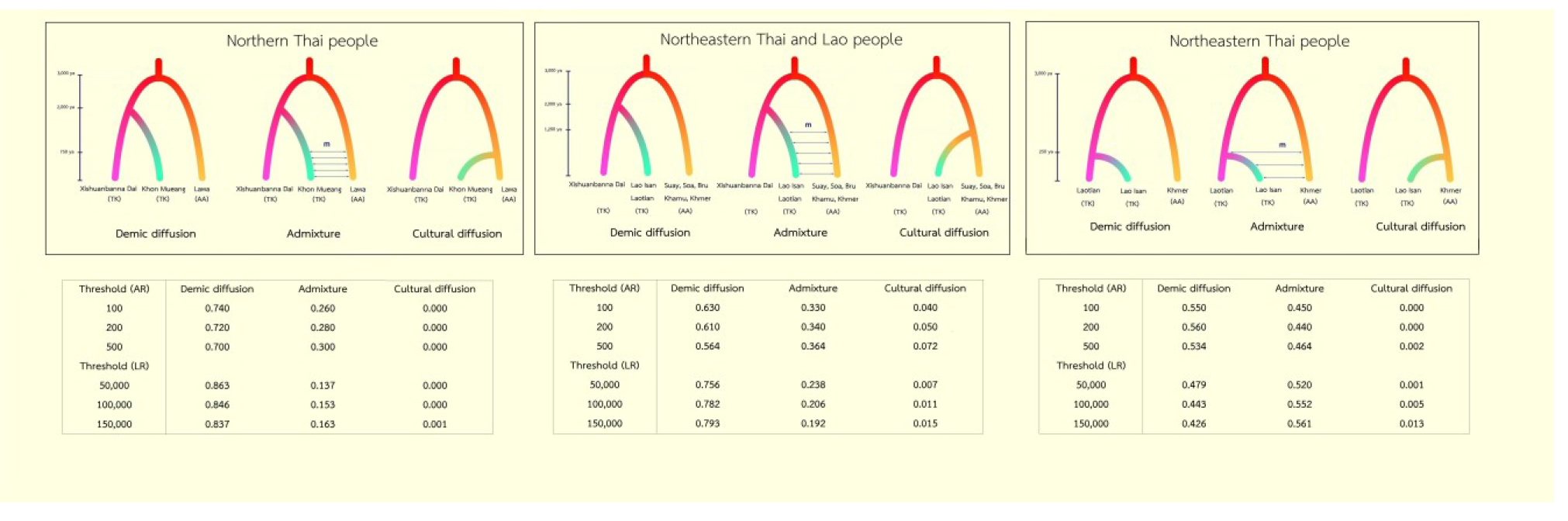
Proposed demographic models for three independent ABC tests concerning northern Thais, northeastern Thais combined with Laotian, and northeastern Thais. Each test consists of three scenarios according to three hypotheses, i.e. demic diffusion, admixture and cultural diffusion. The tables under each model are posterior probabilities computed by the acceptance-rejection procedure (AR) and by the weighted multinomial logistic regression (LR) approaches.

Although these three demographic scenarios are proposed for all TK people, archaeological, linguistic and historical evidence clearly indicate the potential for differences in the local history and demography, especially for groups from northern vs. northeastern Thailand^10,33^. We therefore performed the analyses of Approximate Bayesian Computation (ABC) using three different data sets in all three demographic scenarios: (1) northern Thai people (Khon Mueang, KM); (2) ethnic Lao including northeastern Thai people (Lao Isan, IS) and Laotian (LA); and (3) Lao Isan (to infer the history of this specific population, for reasons detailed in the Methods section). In each analysis, we used AA populations for comparison and set priors for some parameters (e.g. divergence and admixture time) based on historical evidence, as detailed in the Methods section.

In general, the results of the ABC analyses show that in all cases the simulated data included the observed data (Supplementary Fig. 7) and the results of the model selection are consistent among different thresholds, i.e. the different numbers of simulations retained to fit the logistic regression curve. The highest posterior probabilities in both approaches, acceptance-rejection procedure (AR) (0.70-0.74) and weighted multinomial logistic regression (LR) (0.84-0.86), support the demic diffusion model in the northern Thai KM (Fig. 7). Even though the AA-speaking LW groups have culturally interacted with the KM^9–10^, they are not the maternal ancestor of the KM. The test of ethnic Lao (IS and LA; scenario 2) shows the same trend in supporting the demic diffusion model, although it received higher support by LR (0.76-0.79) than by AR (0.56-0. 63). The ethnic Lao are thus genetically distinct from the neighboring AA speaking groups, including the KH, KA SO, SU and BU groups. These two results for TK groups across a vast area of Thailand and Laos thus indicate a genetic origin of the TK from southern China followed by a rapid population expansion from (presumably) a few groups to the current census size of around 50 million, within 1 to 2 ky. For the last analysis concerning the origin of the IS population, there is no distinction between the demic diffusion and admixture models, which differ by absence/presence of migration between KH and IS beginning ~250 years ago. The AR assigned a probability of about 0.55 to demic diffusion and about 0.45 to admixture but *vice versa* in LR. In either event, this analysis does not support the purely cultural diffusion model.

The results of power analysis for the three tested datasets indicated that the true positive rate is generally good, in particular for the demic diffusion model in the first two tests (which was unequivocally supported by the model selection procedures). The false positive rate is low in almost all of the comparisons (less than 0.05) for the selected model of the second test, and slightly higher (0.066) for the selected model of the first test (Supplementary Table 5). In sum, these results confirm the reliability of the posterior probabilities of the models.

## Discussion

In conclusion, the extensive and intensive sampling of complete mtDNA genomes in 51 AA and TK groups from Thailand and Laos shows a high genetic diversification with a total of 212 haplogroups observed. The proposed autochthonous ancient lineages are B5a1d, B6a, R22, R9b and F1f; the many basal lineages detected in this study suggests that the area of present-day Thailand and Laos may have been an ancient migratory route for modern humans, in accordance with the finding that the oldest modern human remains in East Asia are from Tam Pa Ling cave in Laos^1^. Previous studies have suggested Myanmar^34^ and Cambodia^26^ as the corridor for initial settlers, assuming travel along river valleys; our results indicate that in addition, early modern human groups may have migrated through the interior upland, as also suggested by archaeological evidence found in caves in the highlands^3,35^.

Several lines of evidence point to a more ancient presence of AA groups than of TK groups, including greater genetic heterogeneity and on average older maternal lineages, in keeping with previous studies^24–25,36^. There are also distinct affinities between some AA groups (especially the Mon groups) and South Asia, where AA groups are also found. TK groups are less heterogeneous, tend to show more signs of population expansion, and more genetic affinities with southern Chinese groups and with AN groups. The modeling of different demographic scenarios for different groups of populations further supports a demic diffusion of the ancestors of TK groups from southern China. The genetic affinities between TK and AN groups are in keeping with linguistic affinities between the TK and AN language families^37^ and may be explained by the hypothesis that Austronesians are descended from a migration from northern China that also continued into southern China and MSEA^27^. There are further genetic affinities between MSEA and ISEA, but no sharing of recent sublineages, in keeping with previous studies that suggested a pre-Austronesian migration from MSEA to ISEA^38^.

Finally, a surprising – and sobering – finding of this study is that there is significant genetic heterogeneity among samples from the same ethnolinguistic group from different locations. This results holds for all cases where there was more than one sampling location per ethnolinguistic group (Table 1). It appears that this heterogeneity arises from various sources. In the hill tribes, such as the Lawa and H’tin, isolation and drift due to geography and cultural constraints (e.g., matrilocality) appear to be the major factor. For the lowland populations (MO, KH, IS, KM, and PU) recent gene flow with other groups seems to be the major factor. In any event, a common assumption in studies of genetic history is that different samples from the same ethnolinguistic group should have (more or less) the same history, and therefore one sampling location is assumed to be representative of the entire ethnolinguistic group. However, it would seem that this assumption should be evaluated carefully, especially in cases where ethnolinguistic groups are distributed across a wide geographic area; where feasible, multiple samples should be taken from the same ethnolinguistic group.

## Methods

### Samples

Blood or buccal samples were collected with informed consent from 1,234 unrelated subjects belonging to 51 populations that were classified into 23 ethnolinguistic groups (Fig. 1 and Supplementary Table 2). All groups speak either AA or TK languages and all are from Thailand, with the exception of two populations from Laos. Approvals for human research for this study were obtained from Chiang Mai University, Khon Kaen University, Naruesuan University, and the Ethics Commission of the University of Leipzig Medical Faculty.

### MtDNA sequencing and multiple alignment

DNA was isolated as described previously from blood samples^39^ and from buccal cells with the Gentra Puregene Buccal Cell Kit (Qiagen). Sequencing libraries were constructed using a multiplex protocol for the Illumina Genome Analyzer platform^40^ and were enriched for mtDNA as described previously^41^. Several Illumina platforms and lengths of sequencing reads were employed, with post processing using Illumina software and the Improved Based Identification System^42^. The software MIA^43^, which is implemented in an in-house sequence assembly-analysis pipeline for calling consensus sequences and detecting mtDNA heteroplasmy^44^ was used to map sequencing reads to the revised Cambridge Reference Sequence^45^. Details concerning sequencing results and sequence coverage are provided in Supplementary Fig. 7. A multiple sequence alignment of the sequences and the Reconstructed Sapiens Reference Sequence (RSRS)^46^ was executed by MAFFT 7.271^47^.

### Statistical Analyses

The aligned sequences were assigned haplogroups using Haplogrep^48^ with PhyloTree mtDNA tree build 17^49^. MitoTool^50^ was also used to re-check haplogroup assignments. The software Arlequin 3.5.1.3^51^ was used for the following analyses: measures of genetic diversity (Table 1), pairwise genetic distances (*Φ*_*st*_, pairwise difference), AMOVA and a Mantel test comparing genetic and geographic distances between populations; for the latter, we computed three types of geographic distance, i.e. great circle distance, least cost path distance, and resistance distance. The great circle distance matrix was generated by Geographic Distance Matrix Generator v 1.2.3^52^ and the other two distance matrices were computed by the functions *costDistance* in the package gdistance^53^ and using CIRCUITSCAPE^54^ based on a constructed cost-surface raster, respectively. To create this cost-surface raster, briefly, R 3.2.0 was employed using the function *mosaic* from the package raster^55^ to merge two data, i.e. a 30 second elevation grid generated from the WorldClim database^56^ and vector files containing major rivers in Thailand and Laos obtained from NaturalEarth. Then, a cost-surface raster was reclassified with parameters known to affect human movements^57^, e.g. mountain, terrain and river.

Nonparametric MDS analysis (based on *Φ*_*st*_ values) as well as CA analysis using haplogroup counts were constructed using STATISTICA 10.0 (StatSoft, Inc., USA).

BEAST 1.8 was used to construct BSP by population and MCC trees by haplogroup, based on MCMC analyses. The software jModel test 2.1.7^58^ was employed to choose the most suitable model during creation of the input file of BEAST by BEAUTi v1.8^59^. BSP calculations were conducted with the data partitioned between coding and noncoding regions with respective mutation rates of 1.708 × 10^−8^ and 9.8 83 × 10^−8^^60^. Tracer 1.6 (http://tree.bio.ed.ac.uk/software/tracer) was used to visualize the BSP plot. For the BE and CI of haplogroup coalescent times, the RSRS was employed to root the mtDNA tree. The most probable tree from the BEAST runs was assembled with TreeAnnotator and drawn with FigTree v 1.4.0 (http://tree.bio.ed.ac.uk/software/figtree). In order to check clustering of sequences by haplogroup, median-joining networks without pre-or post-processing steps were constructed by Network 4.11 and visualized in Network publisher 1.3.0.0 (http://www.fluxus-engineering.com). Contour maps are generated by Golden Software Surfer 10.0 (Golden Software Inc., USA).

The newly-generated 1,234 mtDNA sequences were compared with a reference data set comprising 2,129 Asian mtDNA genomes representing 62 populations retrieved from the literature (Supplementary Table 4). NJ tree (based on the *Φ*_*st*_) were generated by MEGA 7^61^.

An ABC procedure was employed to choose the best-supported hypothesis about the maternal origins of the Thai and Laotian populations. Owing to the different local histories specific to each region, three different mtDNA data sets from the TK and AA as well as priori parameters (e.g. divergence times) were used in the simulation process (Fig. 7). As the origin time of prehistorical TK speaking groups is unknown, we employed the existing time of the Tai in southern China of ~3 kya, similar to a previous study^62^. Then some prehistorical TK groups started to separate from their common ancestor with the Chinese Dai from their homeland in southern China and spread southward to the area of present-day Thailand in the last 1 to 2 kya^10–12^. Some TK groups finally reached northern Thailand where LW groups are native inhabitants and founded their kingdom, named Lanna around the end of the 13 ^th^ century A.D.^9^. The KM people, the majority of northern Thai, are either genetically from LW groups or admixed with them, and thus should originate at this time. We, therefore, conduct the first analysis by pooling ten KM populations (KM1-KM10) as well as combing the three AA-speaking Lawa groups (LW1-LW3) and using the Xishuanbanna Dai as a representative of the Tai source from southern China^63^. Although nowadays the IS and LA people constitute the vast majority of populations in northeastern Thailand and Laos, respectively, both of them share ethnic identity and the historical motherland of Lao Isan is in Laos^33^. Allowing for the differences in both routes of migration and times of prehistorical TK-groups, the migration from further north to the area of present-day Lao would have met the KH groups, one of the predominant AA people in SEA, who established the Angorian state around 1.2 kya^5^. In addition, SU, KA, BU and SO are the other AA-groups distributed in the area of present-day Laos whose ancestors could have interacted with TK groups. In the second analysis, therefore, the Xishuanbanna Dai is utilized as the Tai sources while all AA groups (KH1-KH2, SU, KA, BU, and SO) are combined and the TK-speaking Lao groups (LA1-LA2 and IS1-IS4) are pooled. In the last analysis, we focus on the IS, as they are a Lao group who recently migrated to northeastern Thailand, approximately 250 ya; evidence of biculturalism between KH and IS in northeastern Thailand has been recorded^64^. One potential scenario was that the IS (IS1-IS4) diverged from the LA (LA1-LA2) without any genetic contact with the KH (KH1-KH2); a second scenario is that IS did admix with KH after diverging from LA. Although an origin of IS from KH is unlikely, we also investigated this scenario.

The simulated datasets were generated by the software package ABCtoolbox^65^. The posterior probabilities were calculated by employing two different approaches, AR^66^, and LR^67^. The former approach considers only a certain number of “best” simulations, and then simply counts the proportion of those retained simulations that were generated by each investigated model. After a few hundred simulations, an excellent fit with the observed data indicates that this approach is reliable^67^, and therefore, 100, 200 and 500 of the best simulations were used in this analysis. According to the latter approach, a logistic regression is fitted where the model is the categorical dependent variable and the summary statistics are the predictive variables. The regression is local around the vector of observed summary statistics, and at the point equivalent to the observed vector of summary statistics, the probability of each model is estimated. Maximum likelihood was used to evaluate the β coefficients of the regression considering different numbers of retained simulations (50000, 100000 and 150000). The posterior probabilities for each model were calculated by the modified R scripts from <u>http://code.google.com/p/popabc/source/browse/#svn%2Ftrunk%2Fscripts</u>. The following summary statistics were employed: the number of haplotypes, haplotype diversity, total number of segregating sites, number of private segregating sites, Tajima’s D, and mean number of pairwise differences for each population, as well as mean number of differences between pairs of populations and pairwise *Φ*_*st*_. The distribution of simulated data under different models with respect to the observed data was evaluated by a visual inspection of a Principal Component Analysis (PCA) of the best 1,000 (or 5,000) simulations for each model, using the PCA function implemented in the R package FactoMineR^68^.

The power to infer the correct model in all tests was estimated by generating 1,000 pseudo-observed datasets according to each analyzed model, with parameter values randomly chosen from the corresponding prior distribution. These pseudo-observed datasets were examined along with the same ABC framework applied in the model selection (i.e. with logistic regression and 50,000 retained simulations). Three different sets of models were considered separately. For each model, we evaluated the proportion of cases where the true model was correctly chosen (i.e. true positives) as well as the proportion of cases where the model selection procedure assigned the highest support to one of the other two tested models (i.e. false positives), considering a posterior probability threshold of 0.5 to assign the support.

## Acknowledgements

We would like to thank all village chiefs and participants who donated their biological samples. We greatly appreciate the assistance of the following coordinators who assisted in collecting samples: Khamnikone Sipaseuth, Saksuriya Triyarach, Narongdech Mahasirikul, Praweena Maneerattanaroongroj, Suparat Srithawong, Kanokpohn Srithongdeang, Nattapol Poltham and Sukhum Ruangchai. We also thank Roland Schröder, Chiara Barbieri, Leonardo Arias Alvis, Enrico Macholdt and Sandra Oliveira from MPI-EVA for technical assistance and valuable advice. This study was primarily funded by the MPI-EVA and Faculty of Science, Khon Kaen University.

## Author contributions

W.K. and M.S. designed the study; W.K., J.K. M.Sr. and D.K. collected the samples; W.K. generated the data; W.K., S.G. A.B. and M.S. analysed the data; W.K., S.G. and M.S. drafted the manuscript.

